# Protein-protein interaction priors shape biologically coherent latent spaces for causally concordant cross-omic translation

**DOI:** 10.1101/2025.10.13.681970

**Authors:** David Martínez-Enguita, Thomas Hillerton, Julia Åkesson, Rebecka Jörnsten, Mika Gustafsson

## Abstract

Deep learning models routinely compress omics into low-dimensional codes, yet many equally accurate embeddings fail to reflect how cells are wired, which limits explanation and causal reasoning. We present a simple, architecture-agnostic approach to make latent spaces biologically legible: a protein-protein interaction (PPI) prior that softly steers autoencoder units to recruit genes that are proximal on the interactome while discouraging redundant reuse. Applied to large DNA methylation (∼155k) and RNAseq (∼993k) compendia and to knowledge-driven (STRING), structure-predicted (RoseTTAFold2-PPI), and union interactomes, this objective reorganizes methylation latents into compact, non-overlapping network neighborhoods without sacrificing reconstruction accuracy. The resulting units map cleanly onto biological processes such as cell-cycle control, immune signaling, proteostasis, mitochondrial metabolism, and RNA handling, with a limited, hub-enriched overlap that plausibly bridges modules. We then asked whether this structured geometry transfers downstream. Using paired TCGA cohorts spanning 23 cancers, omic translators trained on these embeddings, especially a shared-latent bidirectional model, outperformed full-matrix baselines in biologically concordant directions (methylation to transcription, genomics to methylation and transcription) and, crucially, inherited the mechanistic imprint of the upstream encoder. Analytical sensitivity mapping showed that translators fed PPI-guided embeddings preferentially learned known cancer drivers and enriched hallmark pathways, whereas accuracy-matched models trained on non-constrained embeddings did not. Thus, the prior not only regularizes but passes forward a functional coordinate system that makes subsequent predictors mechanistically aware.

By keeping biology in the loss rather than hard-wiring it into the network, our approach scales to very large cohorts, preserves flexibility for understudied genes, and yields latents that are both performant and interpretable. More broadly, it outlines a practical route to mechanism-anchored representation learning that propagates explanatory structure into downstream tasks, advancing explainable AI for multi-omic analysis and clinical decision support.

## 1 INTRODUCTION

Deep learning has become a mainstay tool for modeling high-dimensional molecular profiles. Among its most effective uses is nonlinear dimensionality reduction, which produces compact, reusable features (i.e., latent embeddings) that can be functionally interpreted and used in downstream tasks. Yet, a central problem remains: the same data can be fit equally well by different latent geometries (Semenova *et al.*, 2023), but only some will be coherent with how the cellular interactome operates. In omics particularly, where signals are composite and context-dependent, models can achieve highly accurate performance by conventional metrics while not necessarily encoding explanations reflecting biological processes (Kang, Ko and Mersha, 2022). However, biologically meaningful representations are essential since they provide the foundation for mechanistic inference, allowing hypothesis generation and guiding experimental design by revealing causal structure in complex molecular systems (Mao *et al.*, 2019) (Dwivedi *et al.*, 2020) (Yuan *et al.*, 2021) (Doncevic and Herrmann, 2023) (Martínez-Enguita *et al.*, 2023). Moreover, the ability to trace prediction to function is a key prerequisite for the adoption of decision-making models in clinical settings, as it increases robustness across cohorts, interventional validity, and trust in subsequent analyses (Fortelny and Bock, 2020) (Elmarakeby *et al.*, 2021) (van Hilten *et al.*, 2024). Together, these considerations reinforce the need for latent constraints that keep learned representations aligned with known biology, enabling models to serve as engines for discovery and clinical support.

To address this, a variety of biologically-informed neural networks (BINNs) have been developed. Pathway-aligned factor models such as PLIER (Mao *et al.*, 2019), its successor PLIERv2 (Subirana-Granés *et al.*, 2025), and its transfer-learning extensions MultiPLIER (Taroni *et al.*, 2019) and methPLIER (Takasawa *et al.*, 2024) extract latent factors that map onto curated pathways and cell programs, improving interpretability across cohorts. Bayesian gene-set latent variable models for single-cell RNASeq data, such as f-scLVM and related tools like *slalom* (Buettner *et al.*, 2017), use pathway annotations to guide factors that explain heterogeneity, often refining gene sets in the process. “Visible” or knowledge-primed networks, including DCell (Ma *et al.*, 2018), an early BINN that organized hidden layers into a hierarchical cell ontology; DrugCell (Kuenzi *et al.*, 2020) and P-Net (Elmarakeby *et al.*, 2021), which extended this concept to drug response and cancer phenotypes using multi-level pathway-guided networks; and KPNNs (Fortelny and Bock, 2020), which constrain connectivity to biological networks, embed mechanistic structure into the network’s topology to yield interpretable predictions. Generative encoders with module-tied decoders, like VEGA (Seninge *et al.*, 2021) and expiMap (Lotfollahi *et al.*, 2023), attach latent representations to gene ontology terms or known gene programs for added interpretability; while others like siVAE (Choi, Li and Quon, 2023) jointly learn cell and feature embeddings that recover gene modules linked to phenotypes. Supervised designs also exploit pathway structure: PASNet (Hao *et al.*, 2018) and PathDNN (Deng *et al.*, 2020) use pathway-guided sparsity and wiring to improve prediction while exposing pathway contributions, while the topic-based model scETM (Zhao *et al.*, 2021) couples a neural encoder with an interpretable linear decoder to yield transferable cell and gene signatures. Frameworks like GenNet (van Hilten *et al.*, 2021) perform genotype-to-phenotype prediction with network connections defined solely by prior biology, enabling interpretation at SNP, gene, and pathway levels. For DNA methylation, designs such as MethylCapsNet and MethylSPWNet (Levy *et al.*, 2021) aggregate CpGs into biologically meaningful capsules (e.g., promoters, CpG-island context) for tumor classification, XAI-AGE (Prosz *et al.*, 2024) incorporates pathway priors for multi-tissue epigenetic age estimation, and network-coherent autoencoders (NCAE) (Martínez-Enguita *et al.*, 2023) instead provide an approach to identify methylation latents consistent with the PPI network and extract pathway-enriched signatures. Collectively, these advances point to a general strategy: shape latent spaces with biological structure so that they reflect pathways, modules, and networks rather than unconstrained mixtures, setting the stage for topology-aware guidance of representation learning. Important questions are still unanswered: how can we design soft constraints that both encodes the network topology in a non ad hoc way and supports data-driven embeddings, and how does this design ultimately improve downstream predictions?

The PPIs encode physical and functional proximities among gene products, which we translate into a differentiable objective that encourages each latent unit to concentrate probability on PPI-proximal genes while simultaneously rewarding broad coverage across the genome. The result is a modular latent space in which distinct dimensions specialize in colocalized and functionally consistent network neighborhoods. We adopt standard autoencoders because they accept such latent-space regularization without changing the architecture or imposing additional probabilistic assumptions, they scale efficiently to very large cohorts, and they support discrete unit specialization that maps naturally onto putative pathways. By contrast, variational autoencoders (VAEs) favor continuous latent factors via KL regularization, attention-based models disperse an *a priori* non-topologically-aware signal across many heads, and graph convolutional architectures that directly incorporate the PPI graph into the model layers tend to diffuse information along edges, which risks globally over-smoothing embeddings. Thus, instead of hard wiring genes to units, we keep the interactome as an external guide, so that eligibility is preserved for understudied or weakly connected genes while still biasing the latent representation towards an interpretable and network-coherent structure.

In this work, we introduce a PPI-guided autoencoder framework that uses knowledge- and data-driven interactomes to impose soft topological constraints on the latent space.

By leveraging the protein–protein interaction (PPI) network as a geometric prior, we extend the concept of knowledge-informed learning and shape the latent space of feed- forward autoencoders. This design reinforces our previous findings (Martínez-Enguita *et al.*, 2023; Dwivedi *et al.*, 2020) of co-localization and generates an embedding that co- localizes over the PPI on demand, while preserving accuracy and yielding modular, biologically coherent representations that enhance downstream analyses such as cross-omics translation. We see that this constraint increases the translational between omics in the causative direction, that is from DNA to RNA and when exploring the translational models for cancer it suggests associations of upstream cancer driver genes and pathways which the reference data-driven approach fails to. In summary, this general approach suggests a path how structural knowledge systematically can be integrated in biological models to gradually increase their mechanistic understanding and yield predictive and explainable AI.

## 2 MATERIALS AND METHODS

### 2.1 Data collection and preprocessing

Human pan-tissue DNA methylation profiles and metadata (n = 135,890) generated on Illumina Infinium HumanMethylation450K and MethylationEPIC BeadChips were downloaded from the EWAS Data Hub repository (https://ngdc.cncb.ac.cn/ewas/datahub; accessed 24 March 2025) (Xiong *et al.*, 2020). Additional whole-blood DNAm samples from the Generation Scotland population cohort (n = 18,857) were included (Smith *et al.*, 2006). Raw intensity data (IDAT) files were converted to beta values, when applicable, using the *minfi* R package (v1.54.1).

Between-sample technical variation at the signal-intensity level was removed with Gaussian mixture quantile normalization (GMQN) (Xiong *et al.*, 2022), followed by within-array beta-mixture quantile normalization (BMIQ) to correct the Type II probe design bias. Quality control (QC) and probe filtering were carried out with the *ChAMP* R package (v2.34.0). We discarded (i) non-CpG probes, (ii) probes containing a single nucleotide polymorphism (SNP), (iii) probes mapping to the X or Y chromosome, (iv) multi-hit probes, and (v) probes not present on both Illumina 450K and EPIC arrays.

Missing beta values were imputed by k-nearest neighbors (k = 10) with the *bnstruct* R package (v1.0.15). After preprocessing, 154,747 DNAm profiles with 384,629 CpG sites each were retained for downstream analyses.

Human gene-level RNA-sequencing expression profiles and associated metadata (n = 993,191) were downloaded from the ARCHS4 portal (https://archs4.org; accessed 23 May 2025) (Lachmann *et al.*, 2018). ARCHS4 aggregates publicly available Gene Expression Omnibus (GEO) and Sequence Read Archive (SRA) FASTQ files and processes them with the pseudo-alignment tool Kallisto, summarizing transcript abundances to gene-level estimated counts. Duplicate Gene Symbols were collapsed by averaging their counts, after which library size normalization was applied by converting counts to counts-per-million (CPM) and taking log1p, yielding log-CPM values. The resulting count matrix comprised 19,576 genes (Gene Symbols, Ensembl release 107 annotation).

Paired multi-omic data for the omic translation task was downloaded from the LinkedOmics portal (https://linkedomics.org; accessed 2 April 2025) (Vasaikar *et al.*, 2018). LinkedOmics compiles profiles from The Cancer Genome Atlas (TCGA) and other public repositories. Samples were selected if they had matching DNAm (CpG- level beta values), mRNA expression (gene-level HiSeq log2 RSEM values; RNAseq by expectation-maximization), and somatic copy number variation (SCNV; gene-level GISTIC2 log2 ratios; genomic identification of significant targets in cancer, v2) profiles. Only tumor types with ≥ 50 such triplets were analyzed, leaving 6,723 samples across 23 cancer types. RNAseq normalized expression values were clipped at 15 and minmax-scaled per gene to the interval [0, 1]. SCNV log ratios were clipped at [-2, 2] and minmax-scaled per gene so that 0 mapped to 0.5. DNAm beta values were processed as described in the methylation QC section. Harmonization of gene identifiers across the three modalities yielded a common feature set of 15,354 unique Gene Symbols. Input features and sample metadata used for model training are provided in **Supplementary Files S1 and S2**.

### 2.2 Protein-protein interaction networks as latent space topological constraints

Standard autoencoder architectures efficiently compress gene-level omic profiles, yet the resulting latent dimensions often lack explicit biological meaning. To encourage biologically coherent embeddings while preserving broad gene coverage, we imposed protein-protein-interaction (PPI)-based topological constraints on the latent space: genes that are proximal in a reference interactome are guided to colocalize under the same hidden units, whereas genes without known contacts are free to occupy separate regions. This strategy promotes network neighborhood structure within the latent space, favoring functionally coherent representations, while ensuring that no gene is excluded solely for lacking annotated interactions.

For this purpose, we investigated two complementary interactomes. STRING v12 (Szklarczyk *et al.*, 2023) represents a knowledge-driven resource that aggregates physical bindings, pathway co-membership, co-expression, text mining, and curated databases. STRING interactions are rich in functional context but inherently biased toward well-studied proteins. We retained high-confidence edges with a combined score ≥ 700, yielding 15,966 unique NCBI Gene identifiers linked by 469,676 bidirectional interactions.

As a data-driven counterpart, we used RoseTTAFold2-PPI (RF2, downloaded 30 October 2024 from https://conglab.swmed.edu/humanPPI/humanPPI_download.html), an entirely de novo structural network that derives interaction probabilities from predicted inter-protein contact maps (Baek, 2022) (Zhang *et al.*, 2024). RF2-PPI therefore captures physically plausible interactions, including those involving understudied or low-abundance proteins that may be absent from literature-curated databases. Edges with probability ≥ 0.5 produced 16,060 unique NCBI Gene identifiers and 335,991 unidirectional interactions.

To maximize gene coverage and interaction diversity, we also evaluated the union of the two resources (STRING ≥ 700 ∪ RF2-PPI ≥ 0.5). The merged graph spans 18,650 unique genes and 559,921 interactions, representing a comprehensive yet reliable scaffold for latent-space guidance. The PPI networks used as latent space topological constraints are available in **Supplementary File S3**.

### 2.3 Training of PPI-constrained autoencoders

We trained three-layer autoencoders (latent size *K ɛ* {32, …, 512}, 1.3M to 395M trainable weights) whose latents are regularized by protein-protein interaction structure. At each update we probed the gene map of the latent layer with a "light-up" branch by feeding the identity matrix *I_K_* through the decoder to yield *F* = *f_l_*_ight-up_(*I_K_*), where each row *f_i_* is the gene-score profile generated by latent unit *i*. We convert *f_i_* to a probability over genes with a tempered softmax:

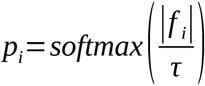

where the temperature τ ɛ [0, ∞) controls the sharpness of the distribution (a lower τ increases sharpness, steering each node towards a small set of high-scoring genes; a higher τ encourages a broader but less discriminative gene allocation). Given a PPI- derived gene-gene distance matrix *D* ɛ *R^N^ ^x^ ^N^*, we encourage each hidden unit to select colocalized genes by minimizing:

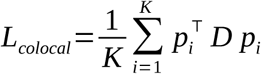

This term encourages each unit to allocate probability mass to PPI-proximal genes, but can drive them toward the same subset. To preserve coverage, we aggregate per-gene maxima across hidden nodes (*m_j_*), representing how strongly each gene is activated by at least one hidden unit. Then, we use a sigmoid (steepness *α,* activation threshold *β*) as a soft indicator for whether gene *j* is covered:

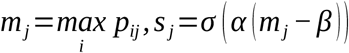

If gene *j* is strongly activated by at least one hidden node, *m_j_* > *β* and *s_j_* ≈ 1 (gene *j* is considered to be covered by the latent space); whereas if gene *j* is not activated strongly enough, then *m_j_* < *β* and *s_j_* ≈ 0 (gene *j* is not considered covered). We reward the union of covered genes as:

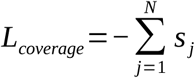

The total objective for PPI-constrained models is thus the weighted sum of standard reconstruction mean squared error (MSE) and the two PPI-aware terms: colocalization loss, penalizing large PPI-graph distances among genes associated with the same hidden unit; and coverage loss, rewarding the inclusion of distinct genes across nodes:

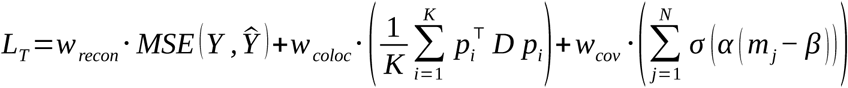

We performed an empirical hyperparameter exploration over a grid of candidate weights and selected the configuration that maximized validation performance. The best trade- off among reconstruction, colocalization, and coverage objectives was obtained with (*w_recon_*, *w_coloc_*, *w_cov_*) = (10, 2.5 x 10^-5^, 3.0 x 10^-7^) and a softmax temperature of τ = 0.1 for all omics; a steepness parameter *α* = 10 and threshold activation *β* = 0.2 for DNAm samples, or *α* = 100 and *β* = 0.01 for RNAseq and SCNV samples. Non-constrained autoencoders (n = 14) were trained with (*w_recon_*, *w_coloc_*, *w_cov_*) = (1, 0, 0). CpG-level latent activations *x_i_*are converted into gene-level activation y_j_ by multiplying the CpG activation vector by a sparse CpG-to-gene mapping matrix *A*. Each non-zero entry of *A* equals 1/n_j_, where n_j_ is the number of CpGs annotated to gene *j*. In practice, *y_j_* is the average of the latent signals for all CpGs *C_j_* mapped to that gene:

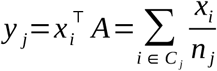

Gene pairs absent from or with no recorded path in the PPI graph are assigned a fixed fallback distance d*_max_* (i.e., colocalization penalty) equivalent to the largest finite distance between any two nodes within the graph. All dense layers used leaky rectified linear unit (leaky ReLU, α = 0.3) activations, and the output was sigmoid-activated. The combined loss was minimized with the Adam optimizer (learning rate = 1 x 10⁻⁴, default β parameters) using mini-batches of 512 or 1,024 samples for DNAm or RNAseq, respectively, with an early stopping patience of 10 epochs. Samples were split into training:validation:test sets with a ratio of 64:16:20.

#### 2.3.1 Weight-slicing adaptation for task-specific fine-tuning of autoencoders

To repurpose DNAm and RNAseq autoencoders pre-trained on large omic compendia for the cross-omic translation task, we applied a weight-slicing strategy that preserves the pre-trained latent manifold while matching task-specific feature sets. The input and output weight matrices were pruned to remove features lacking a counterpart in the translation direction, leaving 242,011 CpG sites or 13,745 unique genes (**Supplementary File S1**). The second and third hidden layers were frozen to preserve the pre-trained latent space (PPI-constrained or non-constrained), whereas the retained rows and columns of the sliced input/output weight matrices were kept trainable, providing a warm start for fine-tuning on TCGA data. These connections were then updated, using the optimizer settings described in Section 2.3, to accommodate cohort-specific distributions without disturbing the biologically-grounded latents learned during pre-training.

#### 2.3.2 Scratch-trained autoencoders for downstream cross-omic translation

For the omic translation task, we trained an additional set of 18 TCGA-only autoencoders from scratch on the intersection of samples available for the relevant omics (DNAm, RNAseq, SCNV; **Supplementary File S2**), using identical train/validation/test splits as the downstream translators. Architecture and optimization follow Section 2.3 (three-layer design, activations, optimizer, early stopping, batch sizes) with the latent bottleneck fixed at 128 dimensions. When the PPI constraint was active (RF2, STRING, Union), we reused the light-up probe and the colocalization and gene-coverage losses as defined in Section 2.3. Non-constrained models used the reconstruction-only objective. For DNAm, CpG-to-gene aggregation followed Section 2.3. No pre-training or weight-slicing was applied. After training, TCGA omic embeddings were exported and used to fit translator models on the train/validation splits. The test set was held out throughout this process, and was used only for final evaluation.

### 2.4 Training of cross-omic translation models

We hypothesized that embedding approaches explicitly enforcing pathway coherence and mutual alignment across omic layers would encourage more biologically meaningful internal representations in downstream models that translate between modalities. To test this, we trained a series of supervised linear and non-linear cross-omic translation models, hereafter referred to as translator models, that learn sample-wise mappings between paired TCGA omic embeddings, and compared their performance when supplied with embeddings generated by PPI-constrained versus non-constrained autoencoders.

We benchmarked four complementary translator model classes: i) deep, fully connected translator neural networks trained on the complete high-dimensional matrices (fullTNN), serving as an upper bound for translation without prior compression; ii) ordinary least-squares (OLS) translators fitted on the encoded embeddings, providing a closed-form baseline for the predictive signal retained by each approach; iii) unidirectional single- hidden-layer translator neural networks trained on embeddings (unidirTNN); and iv) bidirectional shared-latent translator networks (SLTN) that jointly align modality-specific embeddings while optimizing cross-modal and self-reconstruction objectives.

FullTNN models are three-hidden-layer networks of 128 units each (3M to 33M trainable weights, depending on input-output modalities). Every dense layer was followed by batch normalization, leaky ReLU activation (α = 0.3), and dropout (rate = 0.10). A sigmoid-activated output layer matched the [0, 1] scaling of the target features. Models were trained to minimize MSE with the Adam optimizer (learning rate = 1 x 10⁻⁴, default β parameters), using mini-batches of 32 and early stopping after 50 epochs without validation improvement. UnidirTNN models replicated this configuration for a single 128- unit hidden layer (33k trainable weights), with a patience of 1,000 epochs. Linear translators are intercept-free multivariate regressors fitted to translate embeddings from omic A to omic B.

Samples for the translation task were partitioned into training:validation:test sets with a ratio of 64:16:20, and these splits were retained through every autoencoder and translator model training step to prevent data leakage. Each embedding dimension was independently rescaled to [0, 1] using min-max transformation fitted on the training split and applied unchanged to the validation and test sets. To average out performance variability due to model convergence at different optima, we trained four iterations of fullTNNs, unidirTNNs, and SLTNs for every embedding set, for a total of 324 models.

After training, predictions on held-out embeddings were inverted back to the full omic space via the corresponding decoder head, and evaluation was done on the reconstructed complete-data test set using per-feature coefficient of determination (R²), mean absolute error (MAE), mean squared error (MSE), and Pearson correlation. To compare model performance while accounting for dependence across features and iterations, we computed per-feature, per-run R² differences and fit an intercept-only OLS regression, where the intercept estimates the mean ΔR². Inference used two-way cluster-robust standard errors (clusters: feature and run) with small-sample adjustment, and we report the estimate, 95% CI, and P-value. All neural network models were implemented in Keras 2.4.3 running on TensorFlow 2.4.0 (TensorFlow-GPU 2.2.0) under Python 3.8.10. OLS translators and min-max scaling were fitted with *scikit-learn* v1.3.2.

#### 2.4.1 Shared-latent translator networks

Shared-latent translator network (SLTN) models jointly learn a modality-agnostic latent space while performing bidirectional omic imputation and self-reconstruction. Two encoder branches process autoencoder embeddings into latent vectors (Z_A_, Z_B_). From these, four decoder heads are trained in parallel: two cross-modal decoders that predict the opposite omic (A→B, B→A) and two self-reconstruction decoders that rebuild the source omic (A→A, B→B). Each head is formed by a dense layer of 128 units, followed by batch normalization, leaky ReLU (α = 0.3) and dropout (rate = 0.10). The total loss

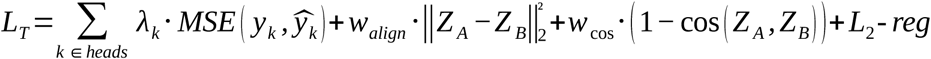

is the weighted sum of: i) bidirectional cross-modal error, expressed as MSE(A, Â_B→A_) + MSE(B, B^_A→B_), as the primary imputation objective; ii) bidirectional self-reconstruction MSE error, MSE(A, A’) + MSE(B, B’), which preserves modality-specific biological signal that would otherwise be sacrificed for cross-modal accuracy; iii) an L2 alignment penalty ||Z_A_ - Z_B_||^2^_2_ that pulls paired latents together sample-wise; and iv) a cosine-similarity term that aligns directions while preventing collapse to a constant vector. We applied the weighting scheme λ_K_ ɛ [w_cross1_ = w_cross2_ = 1.0, w_self1_ = w_self2_ = 0.25], w_align_ = 0.05, w_cos_ = 0.1. Every dense layer was regularized with L2 weight decay (λ = 5 x 10⁻⁷) to smooth the mapping, and weights were initialized using He uniform initialization. The combined loss was minimized with Adam (learning rate = 1 x 10⁻⁴, default β parameters), using mini- batches of 32 and early stopping patience of 1,000 epochs. After training (∼133k trainable weights), either encoder (or both) can project input data into the shared space, from which the corresponding decoder recreates the missing modality, preserving the biologically-meaningful latent structure.

#### 2.4.2 Latent-importance-based feature extraction via analytical sensitivity mapping

We estimated feature-level importance per omic by computing the analytical Jacobian of each SLTN translation path, from the cross-modal prediction layer to the corresponding autoencoder decoder, yielding the exact partial derivatives of each output feature with respect to each latent unit. These derivatives were averaged across samples to obtain the batch-mean sensitivity of genes to latents. We then weighted these sensitivities by the mean absolute latent activations across samples, producing latent-importance values per feature for ranking.

For each paired TCGA sample (*A*, *B*) in the merged training and validation sets of the translation task, we first passed both omics through their fine-tuned autoencoders up to the 128-dimensional third hidden layer. The resulting embeddings were minmax-scaled and fed into the pre-trained, best-performing SLTN encoder branches to extract encoder latent activations. Latent importance (*i*_A_, *i*_B_) was calculated as the sample-averaged absolute activation per unit:

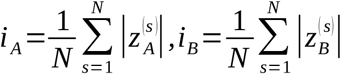

where z ^(s)^ and z ^(s)^ are the latent activation vectors from SLTN encoder branches for sample *s*. To quantify how each latent influences decoded gene outputs, we derived closed-form Jacobians of the translation-decoding map and averaged them across samples:

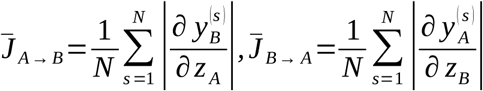

where y ^(s)^ and y ^(s)^ are decoder outputs in gene space. Feature-level importance values were then obtained by projecting latent importance through the element-wise absolute Jacobians:

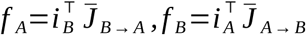

For genes associated with multiple CpG sites, CpG-level importance scores were collapsed to gene-level by retaining the maximum score across all mapped CpGs. For downstream enrichment analyses, we defined the top-activated genes as the 10^3^ highest-ranked features in each list.

### 2.5 Gene annotation and functional enrichment analyses

CpG probes were mapped to genes using the Infinium MethylationEPIC v1.0 B4 Manifest File and probe annotation files via the *ChAMPdata* (v2.40.0) R package. Gene Symbols and Ensembl gene IDs were annotated to NCBI (National Center for Biotechnology Information) gene IDs using *AnnotationDbi* (v1.66.0), *biomaRt* (v2.60.1), and *org.Hs.eg.db* (v3.19.1) R packages.

Human essential genes were retrieved from the Online GEne Essentiality database (OGEE, version 3.09; accessed 5 July 2025) (Gurumayum *et al.*, 2021). We filtered for core-essential genes (CEGs), identified as essential in ≥ 80% of tested cell lines (**Supplementary File S4**). Network-level hub genes were obtained with *networkx* v2.5.1 by ranking PPI proteins by unweighted degree and selecting the top 5% of nodes.

Pleiotropic genes were determined with *mygene* v3.2.2 by querying Gene Ontology (GO) terms from the Biological Process (BP) category for every PPI node. We labeled a gene as pleiotropic when it was associated with ≥ 40 unique GO-BP terms (upper decile of GO annotation distribution). Genes with strong intolerance to loss-of-function (LoF) variation were downloaded from gnomAD v4.1 (Genome Aggregation Database, accessed 6 July 2025) (Chen *et al.*, 2024). A gene was classified as LoF-intolerant when its loss-of-function observed-over-expected upper-bound fraction (LOEUF) was ≤ 0.35, meaning it carries fewer than 35% of the LoF variants expected under neutrality (**Supplementary File S4**).

Functional gene enrichment analysis in KEGG pathways and GO-BP terms was performed using *clusterProfiler* R package (v4.12.6) with default parameters. Hits with FDR-adj. P < 0.05 were considered significantly enriched. Top representative GO-BP terms per hidden node (Figure 4) were selected based on adj. P-value and count ratio of the top 100 ranked genes. Functional relationships between GO terms were inferred by computing the semantic score similarity matrix among top terms across nodes using the *rrvgo* (v1.20.0) R package ("Relevance" distance method), retaining edges with a similarity threshold τ ≥ 0.4. Functional clusters were manually annotated by grouping proximal GO terms with related biological roles into overarching thematic categories.

Reference sets of cancer-associated genes were downloaded (6 August 2025) from the following online resources: 1) Cancer Gene Census (CGC, n = 758) from COSMIC (Catalogue of Somatic Mutations in Cancer) (Sondka *et al.*, 2018), 2) OncoKB™ Cancer Gene List (released 28 July 2025, n = 1,196) (Chakravarty *et al.*, 2017) (Suehnholz *et al.*, 2024), 3) IntOGen (Interactive Onco Genomics) high-confidence drivers (released 20 September 2024, n = 633) (Martínez-Jiménez *et al.*, 2020), and 4) NCG7.2 (Network of Cancer Genes and Healthy Drivers, n = 3,347) (Dressler *et al.*, 2022) (**Supplementary File S5**). We assessed the significance of SLTN top-activated gene enrichment in cancer-associated sets by comparing observed overlaps to an empirical null distribution (k = 10^5^ permutations) generated from the corresponding omic- and PPI- specific background gene universes, mitigating potential inflation of enrichment estimates due to assay design biases.

### 2.6 Statistical analyses and figure editing

Statistical analyses and data processing were conducted in R 4.5.0, within RStudio 2024.12.1, and Python 3.8.10. The threshold for significance corresponds to P-value < 0.05, unless otherwise stated. Figures were created using *ggplot2* v3.5.2, *ggpubr* (v0.6), *ggraph* (v2.2.1) R packages, and *seaborn* (v0.11.1) and *matplotlib* (v3.4.2) Python libraries. Post-processing and final layout adjustments were done in Inkscape v0.92.

## 3 RESULTS

### 3.1 PPI-guided objectives impose network-aware structure on autoencoder latent spaces

First, we asked whether the internal representation learned by autoencoders from large omics compendia is structurally consistent with the human protein-protein interaction (PPI) network, and whether steering this representation improves biological coherence without sacrificing accuracy.

For this purpose, we trained three-hidden-layered, single-omic autoencoders with between 32–512 nodes per layer on DNAm samples from EWAS Data Hub and Generation Scotland (n_A_ ≈ 155,000), and RNAseq samples from ARCHS4 (n_B_ ≈ 993,000). To quantify structure, we identified, for each hidden node, the most strongly activated genes (or CpG-associated genes) and measured (i) their colocalization (harmonic average distance, HAD) on the STRING high-confidence PPI network (score ≥ 700, ∼470,000 interactions) and (ii) the breadth of genes represented across nodes (“coverage”). We then, besides the reconstruction objective, imposed a differentiable constraint on the third hidden layer encouraging each node to learn a compact PPI subgraph (softmax-weighted colocalization loss, Methods) while, in parallel, discouraging redundant use of the same genes across nodes (coverage loss).

Trained RNAseq autoencoders already concentrated their top 1,000 activated genes in nearby regions of the PPI, and did so independently of width and across hidden layers L1–L3 (**Fig. 1a**, green). Average colocalization without constraint ranged between 2.97–3.50 HAD across layers, indicating a strong intrinsic tendency to prioritize interacting genes. In contrast, CpG-mapped genes prioritized by DNAm autoencoders showed little or no PPI colocalization at any width (**Fig. 1a**, red). Average colocalization without constraint was 3.23–4.26 HAD across layers, close to the network baseline and markedly above RNAseq.

**Figure 1.**
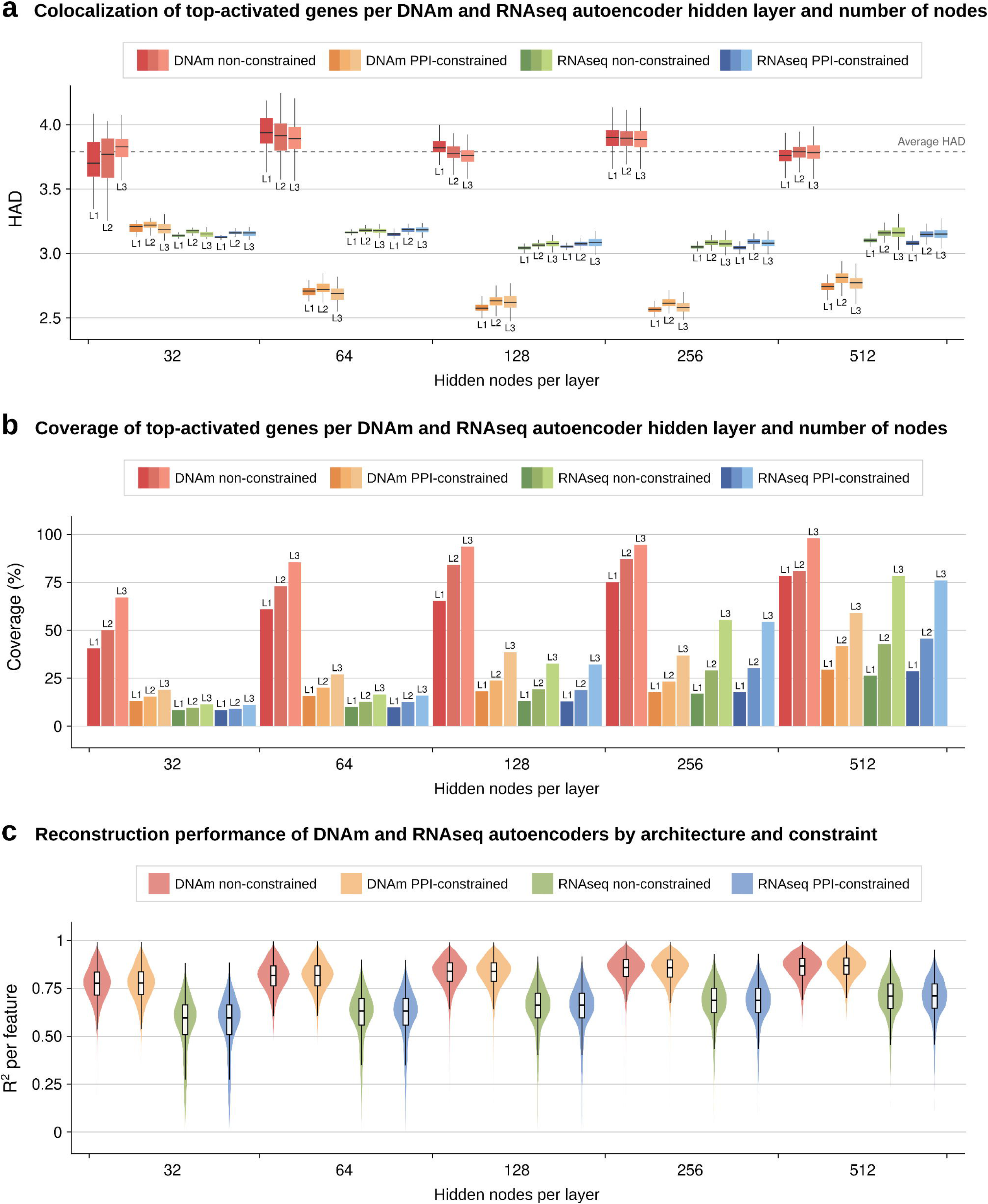
Evaluation metrics for PPI-constrained versus non-constrained autoencoders of 32 to 512 hidden nodes per layer trained using DNA methylation **or transcriptomic data. a)** Colocalization of top-activated genes per autoencoder latent space, measured as inter-gene harmonic average distance (HAD) in the STRING PPI network (interaction confidence ≥ 700) for every hidden node, categorized by hidden layer number and width. The average HAD across all STRING v12 genes (HAD = 3.79) is indicated with a dashed line. **b)** Coverage of top-activated genes per autoencoder latent space, measured as the percentage of unique genes or CpG-associated genes above the activation threshold in at least one hidden node out of the total number of genes, categorized by hidden layer number and width. **c)** Reconstruction performance on the test set for autoencoders per omic and constraint, and hidden layer number and width, measured as the coefficient of determination (R^2^) per output feature.

We next trained models with the PPI constraint acting on the third hidden layer. For RNAseq, colocalization remained essentially unchanged (average with constraint 2.99– 3.45 HAD; **Fig. 1a**, blue), consistent with an already structured latent space. For DNAm, however, the constraint sharply increased PPI coherence: average colocalization rose to 2.47–3.31 HAD across layers (**Fig. 1a**, orange), demonstrating that DNAm representations can be steered to adopt network-aware structure.

The effects of the PPI constraint were most pronounced at intermediate hidden layer widths. DNAm models with 64 to 256 hidden nodes achieved the highest L3 colocalization increases relative to their RNAseq counterparts (ΔHAD = 0.49 at 64, 0.45 at 128, 0.49 at 256; **Fig. 1a**, orange/blue), while 32-node models matched RNAseq and 512-node models began showing decreasing gains (ΔHAD = 0.39). Coverage increased from L1 to L3 and with increasing latent width (**Fig. 1b**), suggesting progressive feature recruitment across layers and expanded capacity to represent additional PPI subgraphs. DNAm models trained without constraints displayed the highest raw coverage (40.6–97.9% of input CpG-associated genes), consistent with diffuse activations spread across many network-distant genes. Introducing the PPI-guided objectives concentrated activations within compact, network-proximal subsets, yielding reduced coverage (**Fig. 1b**, red/orange). Even so, constrained DNAm models outperformed RNAseq in coverage across 32–128 dimensions, nearly two-fold at 32- dim (18.9%, L3), reaching the highest absolute value at 128 (38.7%, L3). Removing the coverage term from the PPI-guided objective lowered coverage on the 128-dim L3 from 38.7% to 31.5% (-7.2 percentage points, -18.6% relative).

Importantly, enforcing PPI structure did not degrade reconstruction (**Fig. 1c**). Across widths and layers, constrained and non-constrained models exhibited overlapping R²- per-feature distributions (32–512-dim median R² = 77.7–86.6, DNAm; 59.5–71.0, RNAseq), indicating that multiple, equally accurate solutions exist and that biologically coherent ones are accessible to gradient-based training.

Together, these results indicate that the PPI-based constraint balances specialization (high local colocalization) with distribution (comparatively broad coverage) and performance, and that an appropriate latent width (64–256) maximizes this balance. Considering this balance alongside model complexity, we selected the 128-dimensional architecture for downstream analyses. Overall, guiding latent spaces with PPI-aware objectives enables DNAm autoencoders to reproduce, and at key widths surpass, the network organization that arises naturally in RNAseq models (**Fig. 1a-c**), setting the stage for downstream tasks that benefit from biologically grounded representations.

### 3.2 A hybrid data- and knowledge-driven prior yields balanced, high-performing latent organization

Having fixed width at 128 hidden nodes, we examined whether the PPI-aware objective generalized beyond STRING to (i) a purely data-driven network (RoseTTAFold2, RF2- PPI) and (ii) the union of RF2 and STRING (Union-PPI). RF2 is a structure-based network inferring interaction probabilities directly from predicted inter-protein contacts (Baek, 2022). We retained RF2 edges with predicted interaction probability ≥ 0.5, resulting in ∼336,000 protein pairs.

We observed that average colocalization was higher (i.e., lower HAD between top- activated genes) on RF2 than on STRING (**Fig. 2a**, left/center), yet the overall trend across models and layers was preserved. RNAseq models showed little change with or without the PPI constraint, as seen previously (avg. HAD L1–L3 without constraint = 2.84; with constraint 2.83–2.84). DNAm models, however, gained clear PPI coherence under the constraint: average HAD decreased from 3.01–3.02 to 2.22–2.25. Coverage trends mirrored those in STRING (**Fig. 2b**, left/center), as coverage increased from L1 to L3 for all models, and PPI-constrained DNAm latent spaces exceeded RNAseq ones (both constrained and not) at L3 by 20.8 and 15.9 pp (+66.3% and +43.9% relative), respectively, balancing PPI focus with breadth.

**Figure 2.**
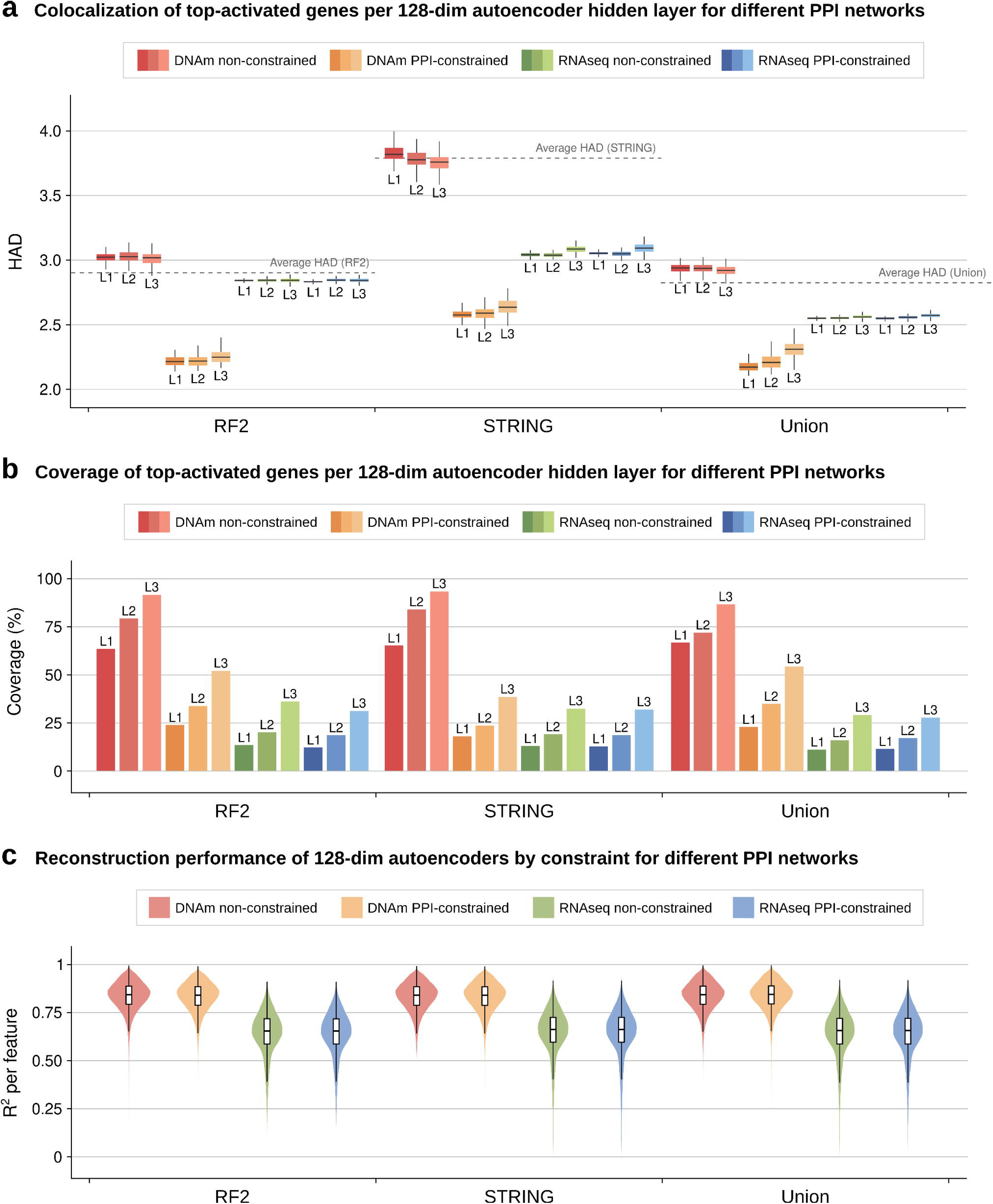
Evaluation metrics for data- or knowledge-driven-PPI-constrained versus non-constrained autoencoders of 128 hidden nodes per layer trained using DNA methylation or transcriptomic data. **a)** Colocalization of top-activated genes per autoencoder latent space, measured as inter-gene harmonic average distance (HAD) in each PPI network for every hidden node, categorized by hidden layer number and PPI network. The average HAD across all RF2, STRING v12, and Union genes (HAD = 2.90, 3.79, 2.82, respectively) are indicated with a dashed line. **b)** Coverage of top-activated genes per autoencoder latent space, measured as the percentage of unique genes or CpG-associated genes above the activation threshold in at least one hidden node out of the total number of genes, categorized by hidden layer number and PPI network. **c)** Reconstruction performance on the test set for autoencoders per omic, constraint, hidden layer number, and PPI network; measured as the coefficient of determination (R^2^) per output feature.

Using the Union network (STRING ≥ 700 ∪ RF2 ≥ 0.5; ∼560,000 interactions) further reduced average HAD (**Fig. 2a**, right), with an evident decline for RNAseq baselines (avg. HAD L1–L3 without constraint = 2.55–2.56; with constraint 2.55–2.57). DNAm models again recovered network coherence under the constraint, reaching HAD levels comparable to RF2 (2.22–2.25 versus 2.18–2.31). Coverage patterns remained robust (**Fig. 2b**, right), with DNAm PPI-constrained autoencoders achieving the highest comprehensiveness (23.0–54.5%, L1–L3), surpassing constrained and non-constrained RNAseq models (26.6 and 25.2 pp, +95.5% and +86.2%, respectively) and exceeding its own coverage on RF2 and STRING by 2.3 and 15.8 pp at L3.

Reconstruction performance was unaffected by the constraint on any network (**Fig. 2c**). R²-per-feature distributions again overlapped within each PPI, showing performance parity for constrained and non-constrained models (128-dim RF2 median R² = 83.9, 84.2 for DNAm; 65.4, 65.4 for RNAseq; STRING median R² = 83.8, 83.9; 66.2, 66.2; Union median R² = 84.3, 84.2; 65.6, 65.6), respectively.

Thus, the PPI constraint generalizes across knowledge- and data-driven interactomes and remains effective on their union. Across networks, biologically meaningful colocalization clustered around HAD ≈ 2.2-2.6, consistent with compact yet non-redundant organization and suggesting an “ideal zone” that balances locality with diversity. Notably, on the Union graph, PPI-constrained DNAm models achieved a nearly two-fold higher coverage at each layer versus RNAseq, while also attaining higher colocalization and reconstruction performance.

### 3.3 PPI constraints separate latent nodes into interactome communities over a shared hub-bridge core

To determine whether the PPI constraint forces each latent feature to capture a densely connected but globally distinct region of the interactome, we calculated the separation score *s_AB_* for every pair of hidden nodes (Menche *et al.*, 2015), using the top 100 most strongly activated genes per node (**Fig. 3a, Supplementary File S6**). A positive value of *s_AB_* indicates that two gene sets are farther apart in the PPI graph than the internal shortest distances of either set, whereas a negative value indicates partial or complete overlap. A usual pragmatic threshold for non-independent gene set pairs is *s_AB_* < -0.01.

**Figure 3.**
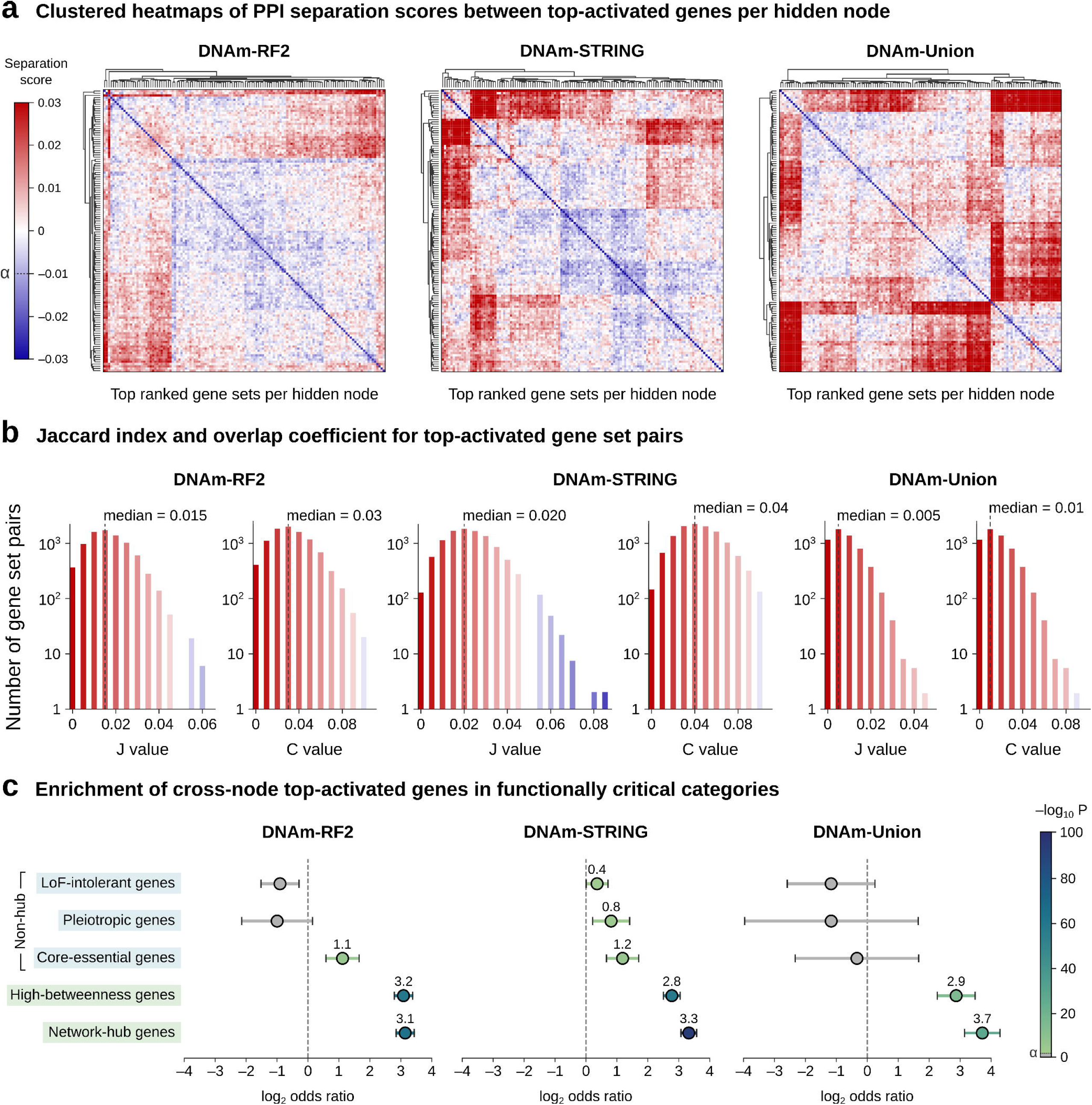
Network separation and similarity metrics for top-activated genes of PPI-constrained, 128-dimensional autoencoders trained on DNAm data. **a)** Heatmaps of PPI network separation scores between top-activated genes per third-layer hidden node. A high separation score (s_AB_ > 0, in red) for a gene set pair indicates that their genes are located farther apart in the PPI graph than within themselves, whereas a low score (s_AB_ < 0, in blue) indicates that a certain overlap exists. The pragmatic threshold for non-independent gene set pairs (*s_AB_* < -0.01) is marked with a dashed line. **b)** Jaccard indices (J) and overlap coefficients (C) for top-activated gene set pairs across PPI-constrained autoencoders. Median values are indicated with dashed lines. **c)** Enrichment of top-activated genes (log2 OR and Fisher’s exact P) shared across different nodes in functionally critical categories, including non-hub known loss-of-function (LoF) intolerant, pleiotropic, and core-essential genes; and high- betweenness, and hub genes.

The distribution of off-diagonal scores across all three interactomes was shifted toward positive values, confirming that the PPI constraint generally kept prioritized latent gene sets apart. The extent of this separation, however, scaled with network completeness. In the STRING network, the median *s_AB_* was 0.0019, with 57.9% of node pairs positively separated, yet 5.5% still fell below -0.01. RF2 showed a nearly identical profile (median *s_AB_* = 0.0011; 56.1% positive; 3.8% negative). In contrast, the denser Union interactome produced a larger rightward shift (median *s_AB_* = 0.0058), with 74.9% of pairs separated and only 1.3% displaying a strong overlap. The dendrograms display several clusters of hidden nodes that remain closer to each other in the interactome, meaning that a subset of the latent space is not entirely orthogonal, even though the global distribution of separation scores is strongly shifted toward independence.

Pairwise Jaccard (J) and overlap coefficients (C) cluster close to zero in all three interactomes (**Fig. 3b**). For the RF2-constrained model, median values are J = 0.015 and C = 0.03; for STRING, J = 0.02 and C = 0.04; and for the Union network J = 0.005 and C = 0.01. By contrast, two random gene lists drawn from genes present in the PPIs would be expected to score only about 0.0033 (Jaccard) and 0.0065 (overlap). Hence, STRING and RF2 medians are five- to six-fold above chance, whereas the Union medians are 1.5-fold higher. Despite this enrichment, the absolute overlaps remain small, typically two to four genes, or 2–4% of each top-activated gene set, supporting the strong general inter-node separation but indicating that a small subset of genes is reused across latent features.

To investigate why certain genes escape the PPI-level separation constraint, we next catalogued the genes that were strongly activated (top 10^2^) in at least 5% of hidden nodes and analyzed their enrichment in hallmarks of biological indispensability: network hubs, high-betweenness genes; and non-hub core-essential genes (CEGs), functionally pleiotropic regulators, and intolerance to loss-of-function (LoF) variants (**Fig. 3c**). We found that shared subsets were significantly enriched across PPI-constrained models in hubs (OR = 8.9, 10.1, and 13.3; Fisher’s exact P = 2.6 x 10^-63^, 2.8 x 10^-94^, and 6.6 x 10^-27^; for RF2, STRING and Union, respectively) and betweenness-central genes (OR = 8.6, 6.9, and 7.4; P = 4.7 x 10^-60^, 1.1 x 10^-58^, and 7.0 x 10^-14^). Non-hub CEGs were overrepresented in STRING-constrained (OR = 2.36, P = 7.5 x 10^-5^) and RF2- constrained shared genes (OR = 2.17, P = 2.4 x 10^-4^). By contrast, enrichment for non- hub pleiotropic and LoF-intolerant genes was context-dependent, with only STRING- constrained models showing moderate overlaps in both categories.

Taken together, these findings indicate that PPI-constrained autoencoders organize the latent space into compact, network-resolved gene communities, with the strongest inter- node separation on the denser Union interactome. The small subset of genes that recur across hidden nodes is non-random and enriched for network hubs and module- bridging genes, explaining the residual clustering seen in the separation maps.

Consistent with the prior evidence that the Union graph yields the most favorable colocalization-to-coverage ratio for DNAm models, this network-level organization is both separated and broadly traversed. We therefore carry forward the 128-dimensional DNAm autoencoder trained on the Union PPI network as flagship model for downstream analyses.

Given the observed modularity of the Union-constrained DNAm latent space, a key question was whether the PPI-localized regions prioritized by individual hidden nodes correspond to specific biological mechanisms. For each node, we ranked genes by activation in the third hidden layer, performed GO-BP enrichment on the top 100 genes, and designated the most significant term as the node’s principal function. Terms were allowed to repeat across nodes and when a top term fell below the multiple-testing threshold (adj. P < 0.05) we retained it to capture the nearest functional assignment. Across the latent space, 92.1% of top terms (n = 114) were unique to a single node, with the most highly enriched terms overall being "regulation of mitotic metaphase/anaphase transition" in node 16 (adj. P = 1.8 x 10^-4^, enrichment score ES = 3.74), "positive regulation of humoral immune response" in node 88 (adj. P = 2.1 x 10^-4^, ES = 3.68), and "cytoplasmic translation" in node 11 (adj. P = 2.1 x 10^-4^, ES = 3.68) (**Supplementary File S7**).

We then embedded the assigned terms in a functional graph by computing semantic similarity between terms and connecting pairs with similarity τ ≥ 0.4. Communities were annotated by thematic proximity (**Fig. 4**). The map resolved into interpretable clusters spanning major cellular systems, including "Protein production and proteostasis" (15 nodes, 11 terms; dark blue), "Immune and inflammatory processes" (12, 12; purple), "Organelle and membrane remodelling" (9, 8; light blue), "Ribosome and RNA processing" (8, 7; orange), and "Mitochondrial function and bioenergetics" (8, 7; yellow), among others. Node indices were distributed across clusters, indicating no relation between node numbering and function.

Thematically, these communities align with biology shaped by DNA methylation: cell- cycle control and proteostasis during proliferation; immune and inflammatory programs; mitochondrial and metabolic regulation; RNA processing and translation; and receptor- proximal signalling. The PPI-guided objective therefore steers latent features toward methylation-sensitive modules rather than mixed signals. Beyond discrete modules, the semantic network also revealed bridges linking clusters, for example macromolecule localization linking secretion and vesicle traffic to translation and immune signalling, indicating that higher-order coupling between processes was preserved while individual nodes specialized.

In sum, imposing the PPI constraint on the Union interactome organizes the DNAm latent space into compact, functionally coherent communities across a wide range of cellular mechanisms, while permitting limited but biologically meaningful cross-talk through module-bridging themes. Each hidden node thus focuses on a definable function, providing a mechanistic map of the representation learned from large-scale DNAm data.

### 3.4 Embedding-based cross-omic translators outperform full-data baselines in causal directions while maintaining accuracy under PPI constraints

Building on this mechanistic map, we next asked whether the learned structure propagates into downstream tasks. We selected cross-omic translation as the downstream assay since it requires sample-wise alignment across modalities and is sensitive to the geometry of the latent space, thereby directly testing whether interactome-aligned embeddings carry forward biological structure. To investigate this, we used paired TCGA cohorts spanning 23 tumor types with DNAm, RNAseq and SCNV profiles (6,723 samples), harmonized to a common gene set (15,354 unique Gene Symbols and 242,011 gene-associated CpGs). Autoencoders produced 128- dimensional embeddings for each omic, either (i) fine-tuned from models pre-trained in large compendia with weight slicing to preserve the latent manifold, or (ii) trained from scratch on TCGA data only. Both PPI-constrained (Union network) and non-constrained variants were evaluated.

We thus trained and benchmarked four translator families that learn sample-wise mappings between modality pairs: a deep fully connected network on the full matrices (FullTNN, no prior compression), ordinary least squares on embeddings (OLS, closed- form baseline), a single-hidden-layer unidirectional translator on embeddings (unidirTNN), and a bidirectional shared-latent translator network (SLTN) that jointly aligns modality-specific embeddings while optimizing cross-modal and self- reconstruction losses. We evaluated performance on a held-out test set as R² per output feature. To stabilize estimates, each neural translator configuration was trained in four independent runs with fixed train/validation/test splits retained across all autoencoder and translator stages.

Embedding-based translators consistently outperformed the full-matrix baseline in biologically concordant directions (**Fig. 5a-b, Supplementary File S8**). In DNAm→RNAseq, SLTN achieved the highest accuracy with R² = 0.547 and 0.546 for constrained and non-constrained fine-tuned embeddings, respectively, both exceeding FullTNN (0.504) by a significant margin (ΔR²_SLTN_constrained–FullTNN_ = 0.041, 95% CI 0.024–0.058, P = 4.64 × 10⁻³; ΔR²_SLTN_nonconstrained–FullTNN_ = 0.040, 95% CI 0.023–0.058, P = 5.06 × 10⁻³) (**Fig. 5a**, left). The same pattern held for scratch-trained embeddings (SLTN_constrained_ = 0.545, SLTNnon_constrained_ = 0.547 vs FullTNN 0.504; ΔR² = 0.038, 95% CI 0.023–0.054, P = 4.07 × 10⁻³; and 0.041, 95% CI 0.027–0.055, P = 2.68 × 10⁻³) (**Fig. 5b**, left).

**Figure 4.**
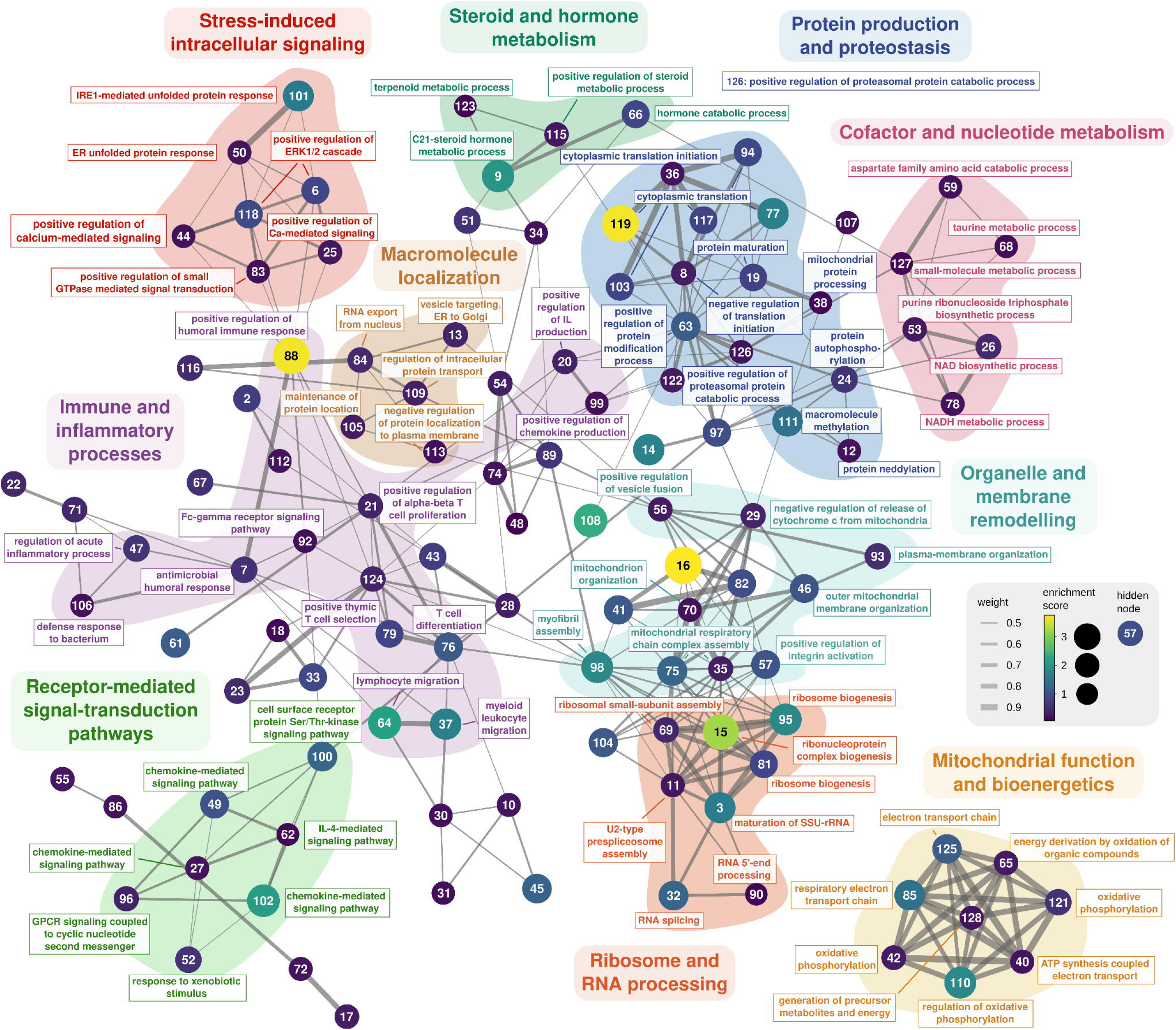
Semantic similarity network of the top enriched GO Biological Process (GO-BP) term per hidden node of the Union-constrained DNA methylation autoencoder (128-dimensional latent space). Each node represents the single most enriched GO-BP term for one hidden node. Labels show the term name and the number inside indicates the hidden-node index. Node size and color represent enrichment significance (-log10 adjusted P) for the selected term. Edges are weighted by semantic similarity, with wider edges indicating higher similarity. Nodes are grouped by graph proximity, and clusters are manually annotated by shared function. Nodes outside the main connected component are omitted.

**Figure 5.**
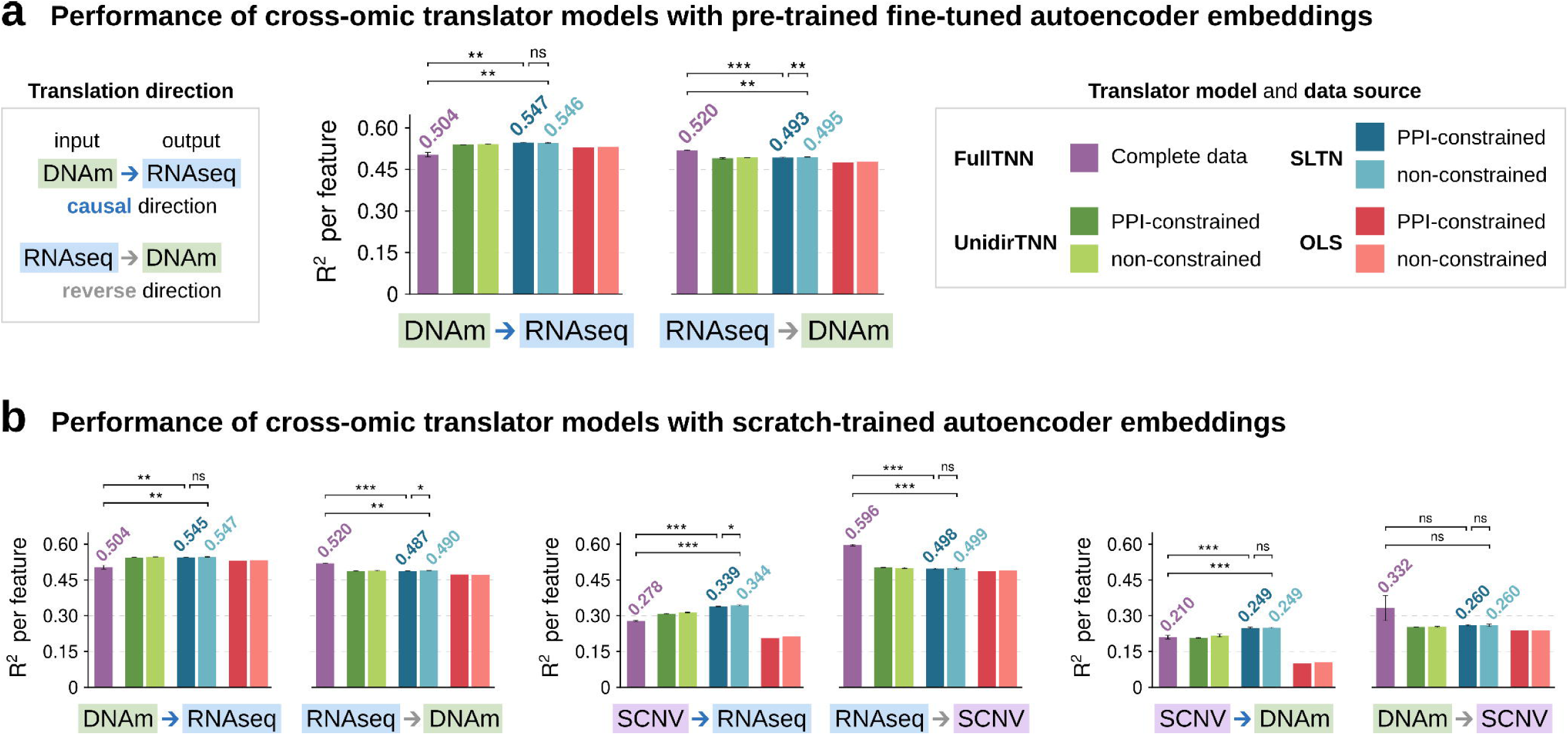
Cross-omic translator performance on paired TCGA cancer sample autoencoder embeddings. Translators trained on the 128-dimensional embeddings from **a)** fine-tuned or **b)** scratch-trained autoencoders, with or without a constrained latent space built on the Union interactome. Models include unidirectional translator neural networks (unidirTNNs), shared-latent translator networks (SLTNs), ordinary least squares (OLS) predictors, and baseline translators trained on full omic matrices (FullTNN). Arrows indicate translation direction; color denotes biological directionality: blue = causally concordant direction. Bars show median coefficient of determination (R²) across output features on the test set, averaged across four independent runs, where applicable. Error bars denote SD across runs. Brackets indicate significance for the average R² shift, estimated via intercept-only OLS on per-feature, per-run differences with two-way cluster-robust standard errors. The test set was withheld throughout both autoencoder and translator training and used only for final evaluation.

The advantage of embedding translators was also evident in SCNV→RNAseq and SCNV→DNAm (scratch-trained). For SCNV→RNAseq, SLTN reached 0.339 (constrained) and 0.344 (non-constrained) versus FullTNN 0.278 (ΔR² = 0.058, 95% CI 0.052–0.064, P = 7.53 × 10⁻⁵; and 0.062, 95% CI 0.052–0.073, P = 2.81 × 10⁻⁴) (**Fig. 5b**, center). For SCNV→DNAm, SLTN attained 0.249 for both constrained and non- constrained embeddings versus 0.210 for FullTNN (ΔR² = 0.053, 95% CI 0.043–0.063, P = 4.71 × 10⁻⁴; and 0.055, 95% CI 0.042–0.067, P = 8.28 × 10⁻⁴) (**Fig. 5b**, right).

Across directions, SLTN delivered the highest absolute R² among embedding-based translators, with unidirTNN performing moderately and OLS lagging behind.

Constrained and non-constrained embeddings produced nearly equivalent accuracy in most settings (e.g. DNAm→RNAseq fine-tuned P = 0.562, ΔR² = 5.89 × 10⁻⁴; scratch-trained P = 0.057, ΔR² = 2.36 × 10⁻³) (**Fig. 5a-b**, left), with a few contrasts such as RNAseq→DNAm and SCNV→RNAseq reaching nominal significance but showing negligible effect sizes (|ΔR²| ≤ 0.005) and lacking consistency across directions (**Fig. 5a**, right, **5b**, left-center).

In summary, embedding-based translators consistently outperform full-matrix baselines in biologically concordant directions (DNAm→RNAseq, SCNV→RNAseq, SCNV→DNAm), indicating that PPI-aligned latents retain the cross-modal signal most relevant for translation. Among translator models, SLTNs with a shared latent space achieved the highest accuracy across all settings. Constrained and non-constrained embeddings yielded near-identical results, with occasional significant differences that were inconsistent across directions and trivial in magnitude. Fine-tuned pre-trained autoencoder embeddings performed on par with scratch-trained task-specific models, demonstrating that latent structures learned under PPI constraints transfer robustly to downstream translation tasks with no meaningful loss of accuracy or power.

### 3.5 PPI-guided embeddings imprint cancer-relevant signal on translator latents

After training and benchmarking the cross-omic translators, we examined whether the learned PPI-aligned structure is propagated to their latents. We analyzed the top-activated genes from the top-performing (highest R^2^) SLTN iterations trained on 128- dimensional embeddings from Union-constrained or unconstrained fine-tuned DNAm and RNAseq autoencoders. Gene rankings were derived by latent-importance projection (**Fig. 6a, Supplementary File S9**): SLTN latent activations were summarized as mean absolute latent activity and analytically projected to gene space via a sample-averaged Jacobian from translator to decoder, yielding per-gene importance scores.

**Figure 6.**
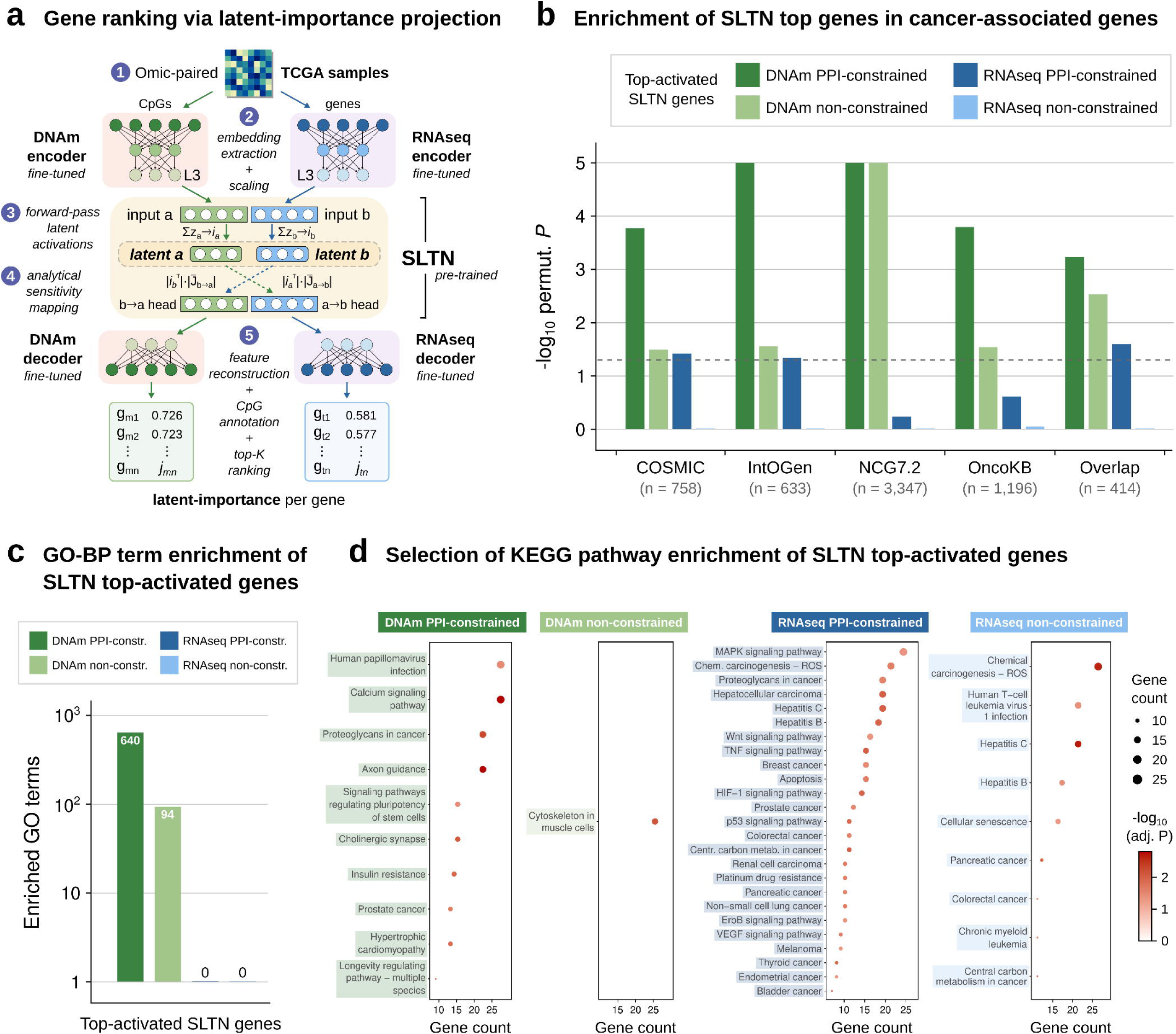
Ranking and functional enrichment of top latent-important genes from SLTN cross-omic translator models. **a)** Schematic representation of pre-trained SLTN gene ranking via latent-importance projection. Paired TCGA omics (1) are encoded with fine-tuned autoencoders to obtain embeddings (2). SLTN latent activations are extracted by forward pass (3) and summarized as mean absolute activation across samples (*i*_a_, *i*_b_; latent importance). These values are then projected to gene space through an analytical, sample-averaged Jacobian from translator to decoder (4), yielding per-gene latent-importance scores used to rank and define top-activated genes (5). **b)** Enrichment in known cancer-associated genes of top latent-important genes from SLTN models trained on embeddings from Union-constrained and non-constrained DNAm and RNAseq autoencoders. P-values were computed from background-specific empirical null distributions (k = 10^5^ permutations). The significance threshold (P = 0.05) is indicated by a dashed line. **c)** Number of significantly enriched (FDR-adjusted P-value < 0.05) GO-BP terms for SLTN top-activated genes per omic and PPI-constraint. **d)** Selection of significantly enriched (FDR-adjusted P-value < 0.05) cancer-relevant KEGG pathways for SLTN top-activated genes per omic and PPI- constraint. Dot size represents the gene count, dot color indicates the enrichment significance.

CpGs were mapped to genes and collapsed so that DNAm- and RNA-derived rankings contain the same set of unique genes, retaining the top 1,000 features from each latent space.

Because translators were trained on paired TCGA tumor samples, we hypothesized that SLTN latents could preferentially rank cancer-associated genes (or their CpG proxies) to support translation, and that this prioritization could be modulated by supplying the biologically coherent PPI-constrained embeddings. For this purpose, we used permutation tests (10⁵ nulls matched to each omic background) for the top-activated gene lists against curated driver resources (COSMIC, IntOGen, NCG7.2, OncoKB, n = 633–3,347 cancer-linked genes) and their intersection (n = 414). We found that DNAm- constrained SLTNs showed the strongest and broadest enrichment: COSMIC (P = 1.70 × 10⁻⁴, 1.55-fold), IntOGen (10⁻⁵, 1.81-fold), NCG7.2 (10⁻⁵, 1.28-fold), OncoKB (1.60 ×

10⁻⁴, 1.42-fold), and the overlap (5.80 × 10⁻⁴, 1.68-fold) (**Fig. 6b**). DNAm-non- constrained SLTNs were also enriched but to a lesser extent (COSMIC 0.032, 1.27-fold; IntOGen 0.028, 1.31-fold; NCG7.2 10⁻⁵, 1.29-fold; OncoKB 0.029, 1.22-fold; overlap 2.91 × 10⁻³, 1.56-fold). RNA-constrained SLTNs exhibited modest enrichment in COSMIC (0.038, 1.26-fold), IntOGen (0.046, 1.27-fold) and the overlap (0.025, 1.39- fold), whereas RNA-unconstrained SLTNs showed no detectable enrichment for any of the lists (**Supplementary File S10**).

Complementing the driver-gene enrichments, functional profiling reinforced the observed pattern. GO-BP analysis of the top-activated genes revealed abundant significant enrichments (FDR-adjusted P-value < 0.05) for DNAm-latent PPI-constrained SLTN top-genes (640 significant terms), reduced but present enrichment for DNAm non-constrained genes (94 terms), and no significant terms for either RNA-derived latent set (**Fig. 6c**). Task-relevant enriched terms for the DNAm-constrained gene set included core oncogenic programs for invasion, survival, stem-like maintenance, and growth-factor-driven proliferation: positive regulation of epithelial-mesenchymal transition (FDR-adjusted P-value = 1.35 x 10^-3^, fold-enrichment FE = 4.23), positive regulation of PI3K-PKB/Akt signal transduction (P = 8.68 x 10^-4^, FE = 2.67), Wnt signaling (canonical/non-canonical; P = 4.52 x 10^-7^ – 0.045, FE = 1.94 – 3.01), TGF-β/SMAD signaling (P = 1.73 x 10^-3^ – 0.031, FE = 1.92 – 3.26), and ERBB/EGFR signaling (P = 0.012– 0.017, FE = 2.31 – 2.34), among others. In comparison, DNAm- non-constrained enrichment included terms such as extracellular matrix organization (P = 0.045, FE = 1.92), small GTPase-mediated signal transduction (P = 2.86 x 10^-3^, FE = 2.45), and macropinocytosis (P = 0.037, FE = 9.29), which are linked to cancer-related programs of matrix remodeling, motility, and nutrient scavenging but overall show a less clear association than the PPI-constrained set enriched terms.

KEGG pathway enrichment corroborated these trends and highlighted disease-relevant programs (**Fig. 6d**). In this case, RNAseq PPI-constrained SLTN top-genes had the strongest signals across canonical oncogenic signaling cascades (MAPK, WNT, mTOR, HIF-1, RAS) and pathways within the KEGG subcategory of Cancer (overview and specific types, 14 hits), including tumor-related processes (chemical carcinogenesis - reactive oxygen species, central carbon metabolism in cancer), and cancer entities (breast, prostate, colorectal, and thyroid cancer, among others), with FDR-adj. P-values between 7.93 x 10^-5^ – 0.047 and fold enrichments (FE) between 1.78–4.80, for a total of 36 significant hits. RNAseq non-constrained latents recovered five pathways within the Cancer subcategory out of 25 significant hits, albeit with attenuated significance (FDR- adj. P between 4.64 x 10^-4^ – 0.049; FE between 1.94–4.60). DNAm-constrained SLTNs were enriched in pathways such as human papillomavirus infection, calcium signalling, proteoglycans in cancer, axon guidance, pluripotency signalling, prostate cancer, and insulin resistance (FDR-adj. P between 1.55 x 10^-3^ – 0.035; FE between 1.91–3.42; 10 significant hits), whereas non-constrained DNAm translator top-activated genes presented no cancer-relevant KEGG enrichment (**Supplementary File S11**).

As a whole, these observations are consistent with the view that training on PPI- constrained embeddings imprints cancer-relevant structure on translator latents, and most strongly for DNAm-based representations. Notably, even where cross-omic translators achieve similar point-estimate accuracy, models trained on inputs guided by a PPI objective exhibit stronger enrichment for cancer-driver genes and associated pathways, indicating that mechanistic organization emerges at a higher representational level under the interactome prior.

## 4 DISCUSSION

We asked whether soft, topology-aware priors based on protein-protein interactions can solve the problematic of equally accurate but biologically divergent latent representations, and whether this alignment to biology transfers downstream to cross-omic reasoning. Four conclusions emerge. First, a PPI-derived objective reliably organizes autoencoder latents into compact and functionally coherent neighborhoods, most clearly for DNAm, where non-constrained models lack intrinsic PPI structure.

Second, this organization generalizes across knowledge-driven (STRING) and structure-predicted (RoseTTAFold2-PPI) interactomes, with their union providing the best balance between local colocalization and genome-wide coverage. Third, embedding-based cross-omic translators increase predictive accuracy specifically in the causally concordant directions (DNAm to RNA, SCNV to RNA/DNAm) relative to the full-matrix baselines, with shared-latent translators achieving the top performance. Fourth, the PPI prior leaves a distinct mechanistic imprint on translator latents trained on constrained autoencoder embeddings.

In line with prior work, latent spaces learned from very large RNAseq compendia naturally concentrate signal on genes proximal in the interactome, consistent with co- expression tracking interaction modules, whereas DNAm representations trained without PPI guidance require careful selection because multiple, equally accurate optima lack biological meaning. Therefore, we introduced a two-part PPI objective: minimizing within-node PPI distances while rewarding non-redundant gene coverage. We observed that this approach reliably pulled DNAm latents toward compact network neighborhoods without degrading reconstruction, indicating that biologically coherent solutions are readily accessible to gradient descent when softly biased by network topology. Practically, this converts diffuse DNAm activations into modular, pathway- aligned representations while keeping standard accuracy metrics intact.

Using either STRING or RoseTTAFold2 PPI networks, and especially their union, yielded consistent gains in latent organization without over-smoothing. The Union graph, which densifies the scaffold to reduce effective distances among true module members while preserving coverage, produced the strongest separation of the gene sets prioritized by each latent node into distinct and functionally independent PPI communities with only limited, biologically plausible overlap. Further, these recurring genes were enriched for hubs and high-betweenness bridges, suggesting a small shared but widespread “core” that connects otherwise specialized modules without undermining overall modularity.

Cross-omic translation use cases confirmed that embedding-based translators do outperform full-data models in the mechanistically concordant directions, with the shared-latent translator network (SLTN) yielding the highest R² across settings. Notably, PPI-constrained and unconstrained embeddings often achieved similar point-estimate accuracy, yet translators trained on PPI-constrained inputs preferentially ranked cancer drivers and hallmark pathways, as quantified by driver-set overlaps and GO/KEGG enrichment, indicating that the prior steers models toward explanations aligned with disease mechanisms without trading off accuracy.

Several limitations to our analysis exist. Current PPI resources are incomplete and context-agnostic, so cell-type- or state-specific interactomes may further sharpen latent specialization. Our colocalization and coverage terms are principled but not unique, as multiscale or hub-penalized distances could also refine the modular separation and reduce trivial overlaps. Additionally, modality-paired perturbation-response data would strengthen the causal claims for our translator models, which rely on observational data. Practically, a mid-sized latent width (128 within a 64 to 256 “ideal zone”), a union interactome prior, and a shared-latent translator provide a strong default. Extending the prior to multilayered regulatory graphs, such as TF-target, kinase-substrate, or ligand-receptor; as well as to single-cell settings, may offer a path to an even more mechanism-aware multi-omic modeling.

In summary, keeping the interactome as a latent-space objective, rather than hard- wiring network topology into the architecture of deep autoencoders, preserved flexibility to recruit weakly connected genes while gently biasing the internal geometry toward biological modules. Regularizing a single, deeper latent layer was sufficient to shape representations without propagating excessive smoothness. Together, these design choices deliver modular and interpretable latents that transfer robustly to downstream tasks, providing a foundation for decision-support AI in clinical contexts.

## Supporting information

Supplementary_Files_all

## ACKNOWLEDGMENTS

Computational resources were granted by the National Academic Infrastructure for Supercomputing in Sweden (NAISS), including National Supercomputer Centre resources Berzelius (Berzelius-2024-5), Sigma (NAISS 2023/5-303), and Tetralith (NAISS 2024/5-385); Chalmers Centre for Computational Science and Engineering (C3SE) resource Alvis (NAISS 2024/5-385); and AIDA Data Hub science platform, part of SciLifeLab Bioinformatics Platform (National Bioinformatics Structure Sweden, NBIS).

## AUTHOR CONTRIBUTIONS

David Martínez-Enguita (Conceptualization [lead], Data Curation [lead], Formal Analysis [lead], Investigation [lead], Methodology [lead], Software [lead], Validation [lead], Visualization [lead], Writing – Original Draft [lead], Writing – Review & Editing [lead]), Thomas Hillerton (Formal Analysis [supporting], Investigation [supporting], Validation [supporting], Writing – Review & Editing [supporting]), Julia Åkesson (Formal Analysis [supporting], Investigation [supporting], Validation [supporting], Writing – Review & Editing [supporting]), Rebecka Jörnsten (Conceptualization [equal], Funding Acquisition [equal], Methodology [supporting], Project Administration [equal], Resources [equal], Supervision [equal], Writing – Review & Editing [supporting]), Mika Gustafsson (Conceptualization [lead], Funding Acquisition [lead], Methodology [supporting], Project Administration [lead], Resources [lead], Supervision [lead], Writing – Review & Editing [supporting]).

## REFERENCES

1. Baek, M. (2022) ‘ Efficient and accurate prediction of protein structures and interactions using RoseTTAFold ’, Acta Crystallographica Section A Foundations and Advances, 78(a1), pp. a235–a235. Available at: 10.1107/s2053273322097649.

2. Buettner, F., Pratanwanich, N., McCarthy, D.J., Marioni, J.C. and Stegle, O. (2017) ‘f- scLVM: Scalable and versatile factor analysis for single-cell RNA-seq’, Genome Biology, 18(1), p. 212. Available at: 10.1186/s13059-017-1334-8.

3. Chakravarty, D., Gao, J., Phillips, S., Kundra, R., Zhang, H., Wang, J., et al. (2017) ‘OncoKB: A Precision Oncology Knowledge Base’, JCO Precision Oncology, (1), pp. 1–16. Available at: 10.1200/po.17.00011.

4. Chen, S., Francioli, L.C., Goodrich, J.K., Collins, R.L., Kanai, M., Wang, Q., et al. (2024) ‘A genomic mutational constraint map using variation in 76,156 human genomes’, Nature, 625(7993), pp. 92–100. Available at: 10.1038/s41586-023-06045-0.

5. Choi, Y., Li, R. and Quon, G. (2023) ‘siVAE: interpretable deep generative models for single-cell transcriptomes’, Genome Biology, 24(1), p. 29. Available at: 10.1186/s13059-023-02850-y.

6. Deng, L., Cai, Y., Zhang, W., Yang, W., Gao, B. and Liu, H. (2020) ‘Pathway-guided deep neural network toward interpretable and predictive modeling of drug sensitivity’, Journal of Chemical Information and Modeling, 60(10), pp. 4497–4505. Available at: 10.1021/acs.jcim.0c00331.

7. Doncevic, D. and Herrmann, C. (2023) ‘Biologically informed variational autoencoders allow predictive modeling of genetic and drug-induced perturbations’, Bioinformatics, 39(6). Available at: 10.1093/bioinformatics/btad387.

8. Dressler, L., Bortolomeazzi, M., Keddar, M.R., Misetic, H., Sartini, G., Acha-Sagredo, A., et al. (2022) ‘Comparative assessment of genes driving cancer and somatic evolution in non-cancer tissues: an update of the Network of Cancer Genes (NCG) resource’, Genome Biology, 23(1), p. 35. Available at: 10.1186/s13059-022-02607-z.

9. Dwivedi, S.K., Tjärnberg, A., Tegnér, J. and Gustafsson, M. (2020) ‘Deriving disease modules from the compressed transcriptional space embedded in a deep autoencoder’, Nature Communications, 11(1), p. 856. Available at: 10.1038/s41467-020-14666-6.

10. Elmarakeby, H.A., Hwang, J., Arafeh, R., Crowdis, J., Gang, S., Liu, D., et al. (2021) ‘Biologically informed deep neural network for prostate cancer discovery’, Nature, 598(7880), pp. 348–352. Available at: 10.1038/s41586-021-03922-4.

11. Fortelny, N. and Bock, C. (2020) ‘Knowledge-primed neural networks enable biologically interpretable deep learning on single-cell sequencing data’, Genome Biology, 21(1), p. 190. Available at: 10.1186/s13059-020-02100-5.

12. Gurumayum, S., Jiang, P., Hao, X., Campos, T.L., Young, N.D., Korhonen, P.K., et al. (2021) ‘OGEE v3: Online GEne Essentiality database with increased coverage of organisms and human cell lines’, Nucleic Acids Research, 49(D1), pp. D998–D1003. Available at: 10.1093/nar/gkaa884.

13. Hao, J., Kim, Y., Kim, T.K. and Kang, M. (2018) ‘PASNet: Pathway-associated sparse deep neural network for prognosis prediction from high-throughput data’, BMC Bioinformatics, 19(1), p. 510. Available at: 10.1186/s12859-018-2500-z.

14. van Hilten, A., Kushner, S.A., Kayser, M., Arfan Ikram, M., Adams, H.H.H., Klaver, C.C.W., et al. (2021) ‘GenNet framework: interpretable deep learning for predicting phenotypes from genetic data’, Communications Biology, 4(1), p. 1094. Available at: 10.1038/s42003-021-02622-z.

15. van Hilten, A., van Rooij, J., Heijmans, B.T., ’t Hoen, P.A.C., Meurs, J. van, Jansen, R., et al. (2024) ‘Phenotype prediction using biologically interpretable neural networks on multi-cohort multi-omics data’, npj Systems Biology and Applications, 10(1), p. 81. Available at: 10.1038/s41540-024-00405-w.

16. Kang, M., Ko, E. and Mersha, T.B. (2022) ‘A roadmap for multi-omics data integration using deep learning’, Briefings in Bioinformatics, 23(1). Available at: 10.1093/bib/bbab454.

17. Kuenzi, B.M., Park, J., Fong, S.H., Sanchez, K.S., Lee, J., Kreisberg, J.F., et al. (2020) ‘Predicting Drug Response and Synergy Using a Deep Learning Model of Human Cancer Cells’, Cancer Cell, 38(5), pp. 672–684.e6. Available at: 10.1016/j.ccell.2020.09.014.

18. Lachmann, A., Torre, D., Keenan, A.B., Jagodnik, K.M., Lee, H.J., Wang, L., et al. (2018) ‘Massive mining of publicly available RNA-seq data from human and mouse’, Nature Communications, 9(1), p. 1366. Available at: 10.1038/s41467-018-03751-6.

19. Levy, J.J., Chen, Y., Azizgolshani, N., Petersen, C.L., Titus, A.J., Moen, E.L., et al. (2021) ‘MethylSPWNet and MethylCapsNet: Biologically Motivated Organization of DNAm Neural Networks, Inspired by Capsule Networks’, npj Systems Biology and Applications, 7(1), p. 33. Available at: 10.1038/s41540-021-00193-7.

20. Lotfollahi, M., Rybakov, S., Hrovatin, K., Hediyeh-zadeh, S., Talavera-López, C., Misharin, A. V., et al. (2023) ‘Biologically informed deep learning to query gene programs in single-cell atlases’, Nature Cell Biology, 25(2), pp. 337–350. Available at: 10.1038/s41556-022-01072-x.

21. Ma, J., Yu, M.K., Fong, S., Ono, K., Sage, E., Demchak, B., et al. (2018) ‘Using deep learning to model the hierarchical structure and function of a cell’, Nature Methods, 15(4), pp. 290–298. Available at: 10.1038/nmeth.4627.

22. Mao, W., Zaslavsky, E., Hartmann, B.M., Sealfon, S.C. and Chikina, M. (2019) ‘Pathway-level information extractor (PLIER) for gene expression data’, Nature Methods, 16(7), pp. 607–610. Available at: 10.1038/s41592-019-0456-1.

23. Martínez-Enguita, D., Dwivedi, S.K., Jörnsten, R. and Gustafsson, M. (2023) ‘NCAE: data-driven representations using a deep network-coherent DNA methylation autoencoder identify robust disease and risk factor signatures’, Briefings in Bioinformatics, 24(5). Available at: 10.1093/bib/bbad293.

24. Martínez-Jiménez, F., Muiños, F., Sentís, I., Deu-Pons, J., Reyes-Salazar, I., Arnedo- Pac, C., et al. (2020) ‘A compendium of mutational cancer driver genes’, Nature Reviews Cancer, 20(10), pp. 555–572. Available at: 10.1038/s41568-020-0290-x.

25. Menche, J., Sharma, A., Kitsak, M., Ghiassian, S.D., Vidal, M., Loscalzo, J., et al. (2015) ‘Uncovering disease-disease relationships through the incomplete interactome’, Science, 347(6224), p. 841. Available at: 10.1126/science.1257601.

26. Prosz, A., Pipek, O., Börcsök, J., Palla, G., Szallasi, Z., Spisak, S., et al. (2024) ‘Biologically informed deep learning for explainable epigenetic clocks’, Scientific Reports, 14(1), p. 1306. Available at: 10.1038/s41598-023-50495-5.

27. Semenova, L., Chen, H., Parr, R. and Rudin, C. (2023) ‘A Path to Simpler Models Starts With Noise’, Advances in Neural Information Processing Systems, 36, pp. 3362–3401.

28. Seninge, L., Anastopoulos, I., Ding, H. and Stuart, J. (2021) ‘VEGA is an interpretable generative model for inferring biological network activity in single-cell transcriptomics’, Nature Communications, 12(1), p. 5684. Available at: 10.1038/s41467-021-26017-0.

29. Smith, B.H., Campbell, H., Blackwood, D., Connell, J., Connor, M., Deary, I.J., et al. (2006) ‘Generation Scotland: The Scottish Family Health Study; a new resource for researching genes and heritability’, BMC Medical Genetics, 7(1), p. 74. Available at: 10.1186/1471-2350-7-74.

30. Sondka, Z., Bamford, S., Cole, C.G., Ward, S.A., Dunham, I. and Forbes, S.A. (2018) ‘The COSMIC Cancer Gene Census: describing genetic dysfunction across all human cancers’, Nature Reviews Cancer, 18(11), pp. 696–705. Available at: 10.1038/s41568-018-0060-1.

31. Subirana-Granés, M., Nandi, S., Zhang, H., Chikina, M. and Pividori, M. (2025) ‘PLIERv2: bigger, better and faster’. Available at: 10.1101/2025.06.05.658122.

32. Suehnholz, S.P., Nissan, M.H., Zhang, H., Kundra, R., Nandakumar, S., Lu, C., et al. (2024) ‘Quantifying the Expanding Landscape of Clinical Actionability for Patients with Cancer’, Cancer Discovery, 14(1), pp. 49–65. Available at: 10.1158/2159-8290.CD-23-0467.

33. Szklarczyk, D., Kirsch, R., Koutrouli, M., Nastou, K., Mehryary, F., Hachilif, R., et al. (2023) ‘The STRING database in 2023: protein-protein association networks and functional enrichment analyses for any sequenced genome of interest’, Nucleic Acids Research, 51(1 D), pp. D638–D646. Available at: 10.1093/nar/gkac1000.

34. Takasawa, K., Asada, K., Kaneko, S., Shiraishi, K., Machino, H., Takahashi, S., et al. (2024) ‘Advances in cancer DNA methylation analysis with methPLIER: use of non-negative matrix factorization and knowledge-based constraints to enhance biological interpretability’, Experimental and Molecular Medicine, 56(3), pp. 646–655. Available at: 10.1038/s12276-024-01173-7.

35. Taroni, J.N., Grayson, P.C., Hu, Q., Eddy, S., Kretzler, M., Merkel, P.A., et al. (2019) ‘MultiPLIER: A Transfer Learning Framework for Transcriptomics Reveals Systemic Features of Rare Disease’, Cell Systems, 8(5), pp. 380–394.e4. Available at: 10.1016/j.cels.2019.04.003.

36. Vasaikar, S. V., Straub, P., Wang, J. and Zhang, B. (2018) ‘LinkedOmics: Analyzing multi-omics data within and across 32 cancer types’, Nucleic Acids Research, 46(D1), pp. D956–D963. Available at: 10.1093/nar/gkx1090.

37. Xiong, Z., Li, M., Yang, F., Ma, Y., Sang, J., Li, R., et al. (2020) ‘EWAS Data Hub: A resource of DNA methylation array data and metadata’, Nucleic Acids Research, 48(D1), pp. D890–D895. Available at: 10.1093/nar/gkz840.

38. Xiong, Z., Li, M., Ma, Y., Li, R. and Bao, Y. (2022) ‘GMQN: A Reference-Based Method for Correcting Batch Effects and Probe Bias in HumanMethylation BeadChip’, Frontiers in Genetics, 12. Available at: 10.3389/fgene.2021.810985.

39. Yuan, B., Shen, C., Luna, A., Korkut, A., Marks, D.S., Ingraham, J., et al. (2021) ‘CellBox: Interpretable Machine Learning for Perturbation Biology with Application to the Design of Cancer Combination Therapy’, Cell Systems, 12(2), pp. 128–140.e4. Available at: 10.1016/j.cels.2020.11.013.

40. Zhang, J., Humphreys, I.R., Pei, J., Kim, J., Choi, C., Yuan, R., et al. (2024) ‘Computing the Human Interactome’, bioRxiv, p. 2024.10.01.615885. Available at: 10.1101/2024.10.01.615885.

41. Zhao, Y., Cai, H., Zhang, Z., Tang, J. and Li, Y. (2021) ‘Learning interpretable cellular and gene signature embeddings from single-cell transcriptomic data’, Nature Communications, 12(1), p. 5261. Available at: 10.1038/s41467-021-25534-2.

